# Cell-free protein synthesis enables one-pot cascade biotransformation in an aqueous-organic biphasic system

**DOI:** 10.1101/2020.03.17.995118

**Authors:** Wan-Qiu Liu, Changzhu Wu, Michael C. Jewett, Jian Li

## Abstract

Biocatalytic cascade reactions have become increasingly important and useful for chemical synthesis. However, biocatalysts are often incompatible with organic solvents, which prohibits many cascade reactions involving nonpolar substrates. In this work, we used cell-free protein synthesis (CFPS) to express enzymes in an aqueous-organic biphasic system for the construction of an artificial enzymatic pathway. CFPS-expressed enzymes without purification performed efficiently to convert styrene (below 20 mM) to (*S*)-1-phenyl-1,2-ethanediol (two steps in one pot) with 100% conversion. In addition, our CFPS system showed great tolerance to different organic solvents and, importantly, the entire biocatalytic system can be consistently scaled up without reduction of the substrate conversion rate. We therefore anticipate that our cell-free approach will make possible cost-effective, high-yielding synthesis of valuable chemicals.

## Main Text

Enzymatic cascades, which have become a green and sustainable method for organic synthesis, provide an interesting approach for manufacturing value-added chemicals and advanced pharmaceutical intermediates (Bornscheuer et al., 2012; Reetz, 2013; Schmermund et al., 2019; Sheldon et al., 2019; Sheldon et al., 2018). Enzymes offer high enantioselectivity, fast conversion rates, and mild reaction conditions. In addition, many natural enzymes have similar reaction conditions in cellular aqueous environments, which are often at nearly neutral pH and ambient temperature. Therefore, enzymes sourced from diverse organisms can be rationally selected to construct natural or artificial enzymatic pathways to conduct chemical transformations amenable for manufacturing (Wu et al., 2018). However, the natural environment for enzymatic biotransformations is in aqueous conditions, which is unfavorable to many industrial reactions with water-insoluble substrates (Klibanov, 2001). To improve the solubility of these types of substrates, water is often replaced with organic solvents as the reaction media but this can denature or reduce catalytic activities of enzymes (Klibanov, 1997). As a result, this made the use of enzymes less attractive for industrial applications. To tackle this problem, aqueous-organic biphasic systems, such as emulsions (Wang et al., 2012; Wiese et al., 2013; Wu et al., 2011) and membrane bioreactors (Boontawan et al., 2006; Doig et al., 1998; Molinari et al., 1997), have been developed for efficient biotransformations. These systems are designed to separate enzymes in a favorable aqueous environment from the organic phase containing the substrate, increasing the aqueous/organic interface and thereby enhancing biocatalytic productivity.

Biotransformations using enzymatic cascades are typically implemented with purified enzymes (*in vitro*), whole-cells (*in vivo*), or even hybrid systems (*in vivo*/*in vitro*) (France et al., 2017). However, each system has its own limitations. For *in vitro* systems, the enzyme purification process is laborious, time-consuming, and costly. For whole-cell biocatalysis, the cell membrane often poses a serious barrier to substrate uptake and product export (Ladkau et al., 2014). Thus, developing new methods to circumvent these obstacles is highly desirable. Recently, cell-free systems have emerged as an attractive and powerful platform for biomanufacturing (Bundy et al., 2018; Carlson et al., 2012; Dudley et al., 2015; Li et al., 2018; Liu et al., 2019; Silverman et al., 2019; Swartz, 2018). In this context, cell-free protein synthesis (CFPS) plays a core role in the production of various proteins *in vitro* (Jaroentomeechai et al., 2018; Kwon et al., 2013; Li et al., 2016; Lu et al., 2014; Martin et al., 2018; McNerney et al., 2019; O’Kane et al., 2019; Wilding et al., 2019). Because of the absence of cell walls, cell-free systems bypass mass transfer limitations and are more tolerant of toxic substrates, intermediates, and products than living microbial cells. In addition, these open cell-free systems allow for easy upstream (*e*.*g*., reaction control and optimization) and downstream (*e*.*g*., sampling and purification) operations. Furthermore, enzymatic reactions can be directly driven via *in situ* CFPS expressed enzymes (biocatalysts) without purification (Goering et al., 2017). However, the potential of CFPS based cell-free systems have not been fully exploited as an efficient approach to carry out cascade biotransformations with water-insoluble substrates.

Here, we use CFPS to enable the one-pot production of (*S*)-1-phenyl-1,2-ethanediol from the nonpolar substrate, styrene, in an aqueous-organic biphasic system (**Figure 1**). We assembled a two-step, artificial enzymatic cascade composed of three enzymes coexpressed in a high-yielding *Escherichia coli* based CFPS system (Li et al., 2016). In the first step, the substrate styrene is converted to an intermediate styrene oxide by a styrene monooxygenase (SMO, which consists of two subunit enzymes StyA and StyB) from *Pseudomonas* sp. strain VLB120 (Panke et al., 1998). In the second step, epoxide hydrolase (SpEH) from *Sphingomonas* sp. HXN-200 (Wu et al., 2013) catalyzes the hydrolysis of styrene oxide to form (*S*)-1-phenyl-1,2-ethanediol, which is a valuable chiral pharmaceutical building block (Bala et al., 2010; Renat et al., 2009). These biocatalytic reactions are carried out in a compartmentalized aqueous-organic emulsion system prepared with a simple block copolymer, poly(ethylene glycol)-block-poly(*ε*-caprolactone (PEG-b-PCL), to provide an exceptionally large interfacial area for efficient biocatalysis (Zhao et al., 2018).

**Figure 1.**
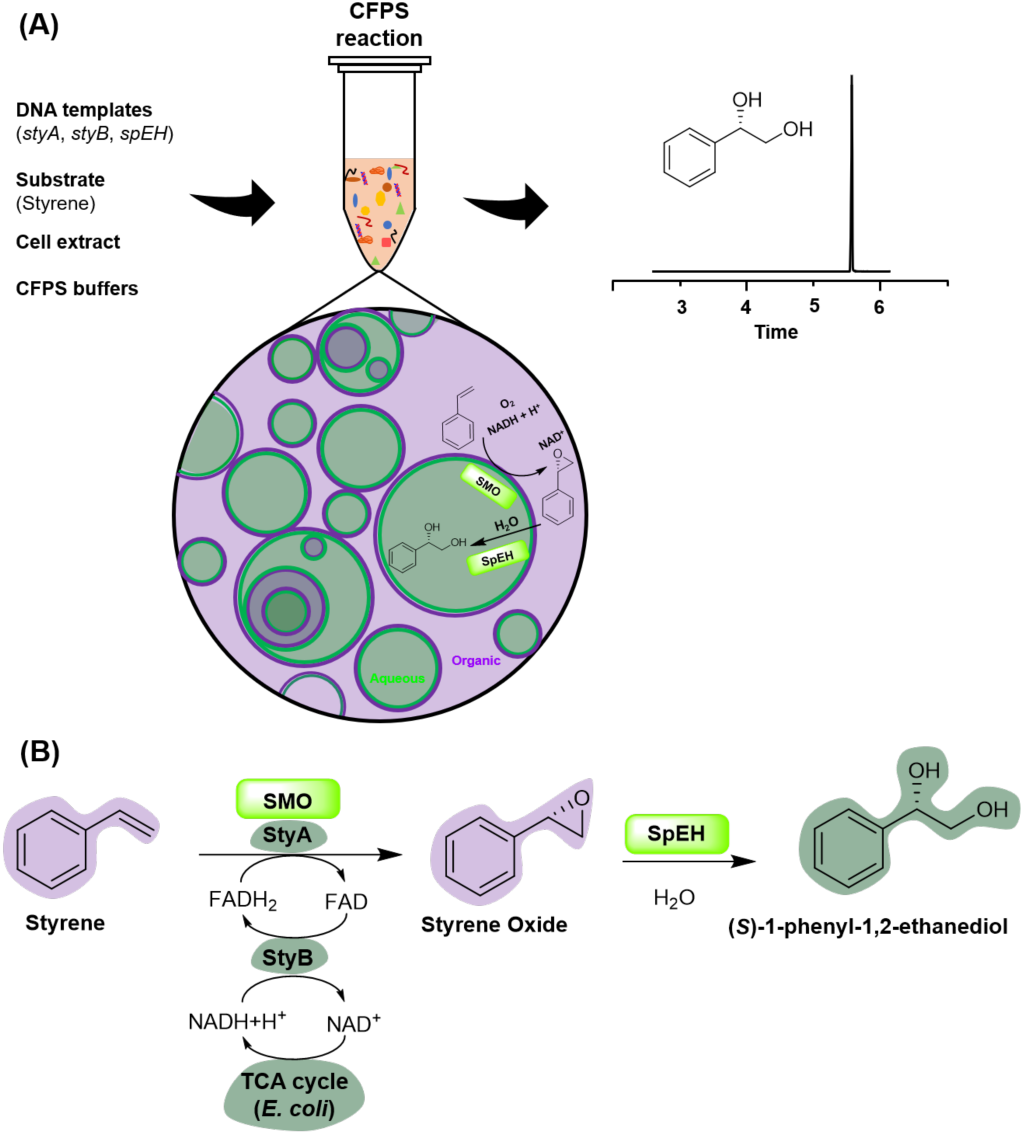
**(A)** Schematic diagram of one-pot cascade biotransformation from styrene to (*S*)-1-phenyl-1,2-ethanediol in a CFPS/emulsion system. SMO: styrene monooxygenase; SpEH: epoxide hydrolase. **(B)** Overall two-step enzymatic cascade reactions. TCA cycle: tricarboxylic acid cycle.

We first determined how active an *E. coli*-based CFPS system is in our aqueous-organic emulsions by expressing super-folder green fluorescent protein (sfGFP). The aqueous CFPS reaction mixture was prepared as reported previously (Li et al., 2016). The organic phase was generated by dissolving 20 mg/mL water-insoluble triblock copolymer (PCL_21_-PEG_45_-PCL_21_) in toluene (Zhao et al., 2018). After mixing the aqueous and organic components at a ratio of 1:1 (v/v), the CFPS reaction was carried out at 30°C and 150 rpm in a shaker for 10 h and, subsequently, the emulsion structure was identified by a confocal laser scanning microscopy (CLSM). As shown in **Figure 2**, both single and multiple compartmentalized structures were observed and sfGFP was synthesized in the aqueous phase with a bright green color. The yield of sfGFP in the aqueous phase reached 1.09 ± 0.01 mg/mL, which is comparable to that in a positive control CFPS reaction (1.33 ± 0.03 mg/mL) without organic phase. These results indicate that our CFPS system is active for protein expression in the aqueous-organic emulsion.

**Figure 2.**
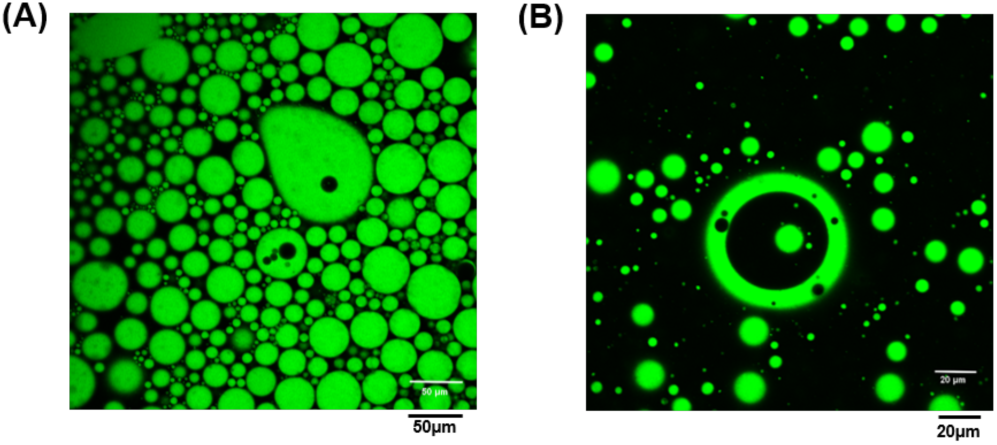
Cell-free synthesis of sfGFP in the CFPS/emulsion system and the single-/multi-compartmentalized structures of the emulsion detected by confocal laser scanning microscopy (CLSM). (**(A)**, 20x and **(B)**, 40x).

We next constructed the artificial enzymatic cascade with StyA (46.3 kDa), StyB (18.4 kDa), and SpEH (42.9 kDa). Prior to one-pot coexpression of these three enzymes, each enzyme was separately expressed by adding 200 ng of plasmid per 15 μL CFPS reaction. Although each enzyme was successfully expressed, their expression levels varied (**Figure 3A**, left). In order to balance the expression level of each enzyme during coexpression, we adjusted plasmid concentrations for each enzyme in CFPS reactions. We found that the best combination of three plasmids was 200 ng (StyA), 50 ng (StyB), and 200 ng (SpEH), according to enzyme expression levels and enzyme solubility (**Figure 3A**, right). In addition, each enzyme was purified (**Figure S1**) and the molar ratio of StyA:StyB:SpEH was about 1:2:1.5 (15.95 μM : 33.70 μM : 23.78 μM). Previous studies reported that the maximal styrene conversion rate was obtained when the molar amount of StyB was equal to or higher than that of StyA (Otto et al., 2004; Panke et al., 1998). Therefore, our CFPS expression system could provide a reasonable enzyme stoichiometry for the enzymatic reactions. Using these conditions, we moved CFPS reactions to the aqueous-organic system for one-pot cascade reactions. To construct the whole biocatalytic pathway, the catalytic ability of each enzyme was initially investigated. The data are shown in **Figure 3B**. Cell-free expression of StyA was sufficient to oxidize styrene (the conversion was not complete) in the absence of StyB, whereas expression of StyB alone did not yield any detectable product of styrene oxide. Interestingly, when StyA and StyB were coexpressed, the activity of StyA for the epoxidation of styrene into styrene oxide was notably restored. Our results agree with previous reports that StyA plays an indispensable role in the catalytic activity of SMO and StyB helps maximize the epoxidation (Otto et al., 2004; Panke et al., 1998). SpEH in our system was also highly active, converting styrene oxide to (*S*)-1-phenyl-1,2-ethanediol without any detectable substrate remaining. While coexpression of StyA and SpEH allows formation of (*S*)-1-phenyl-1,2-ethanediol from styrene at low levels, coexpressing all three enzymes together in one pot enables a higher conversion of styrene to (*S*)-1-phenyl-1,2-ethanediol with no detectable styrene oxide intermediate remaining.

**Figure 3.**
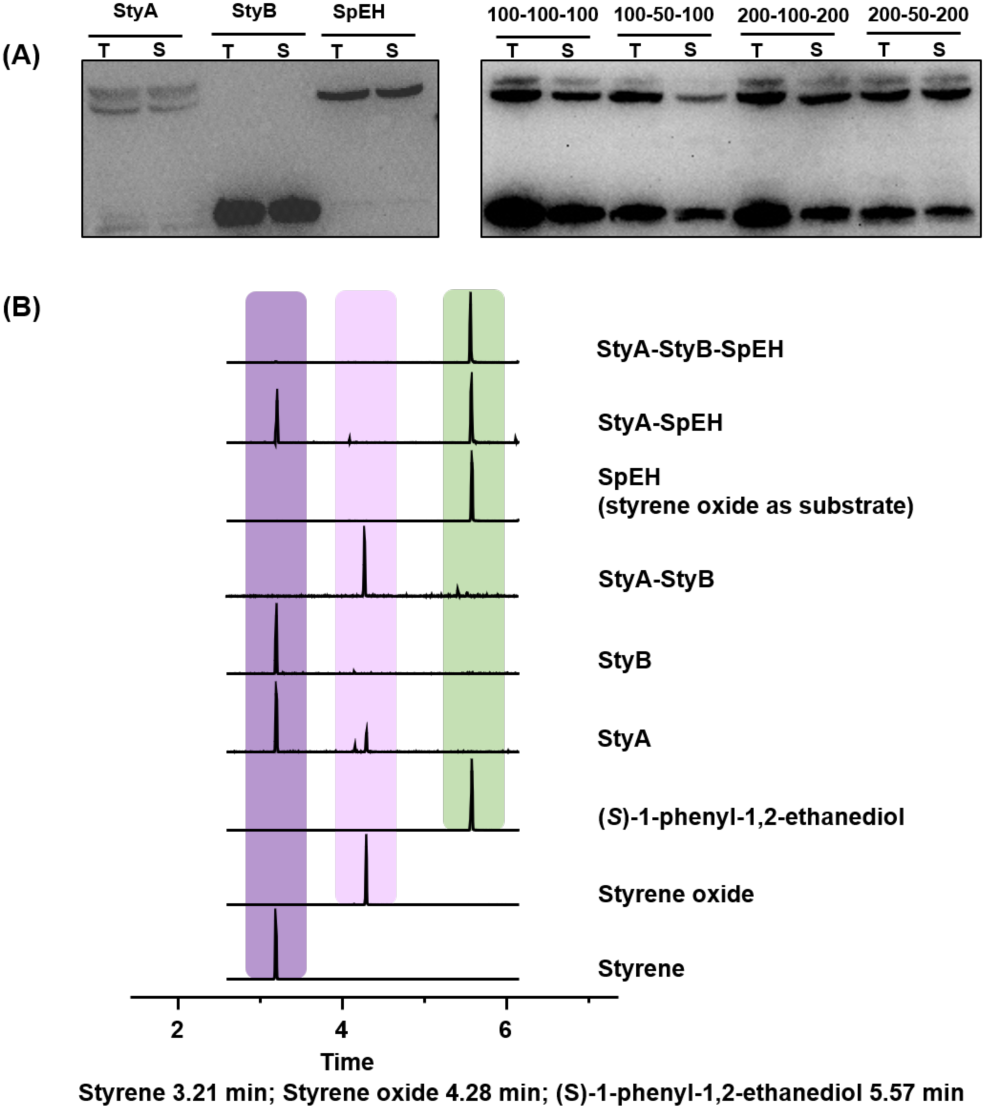
**(A)** Western-blot analysis of individual expression (left) and coexpression (right) of StyA (46.3 kDa), StyB (18.4 kDa), and SpEH (42.9 kDa). T: total protein, S: soluble protein. Numbers on the top of the right panel indicate three plasmid concentrations (StyA-StyB-SpEH) per 15 µL CFPS reaction. Note that the below band of StyA (left) is a truncated protein. **(B)** GC-MS detection of substrate (styrene), intermediate (styrene oxide), and product ((*S*)-1-phenyl-1,2-ethanediol). The concentrations of styrene and styrene oxide were 10 mM when they were used as substrates in the reaction. Three independent reactions were performed for each condition, and an example trace is shown.

Next, we set-out to optimize our system to enhance the cascade productivity. We began our optimization by investigating the concentrations of three plasmids used per 15 μL CFPS reaction. As shown in **Figure 3A**, the optimal plasmid ratio was StyA:StyB:SpEH = 4:1:4 (200-50-200 ng/15 μL), which gives rise to maximum soluble expression of each enzyme in one pot. Therefore, we chose to add different plasmid concentrations to reach the same plasmid ratio. The results indicate that the highest yield of (*S*)-1-phenyl-1,2-ethanediol was achieved with the addition of more plasmids at 200-50-200 ng (**Figure 4A**). However, further increase of plasmids (300-75-300 ng) obviously reduced the product yield. Cell-free reaction temperature is often a key parameter for optimization because it affects enzyme expression and activity (Li et al., 2017). We therefore compared the product yields at different reaction temperatures ranging from 20 to 35°C. Our data suggest that the productivity at 30°C reached the highest level, which is 2-4 times higher than that at other temperatures (**Figure 4B**). Since the optimum reaction temperature in the *E. coli*-based CFPS system is 30°C as reported previously (Goering et al., 2017; Li et al., 2016). we decided to use this temperature in our following cascade biotransformation. After the above primary optimization, although the final product (*S*)-1-phenyl-1,2-ethanediol was produced, the conversion rates based on substrate (10 mM) were still low (maximally about 40%) (**Figure 4B**). We hypothesized that the low expression of StyA in CFPS (**Figures 3A**) likely limited the conversion rate because StyA predominantly contributes to the catalytic activity of SMO (Otto et al., 2004; Panke et al., 1998).

**Figure 4.**
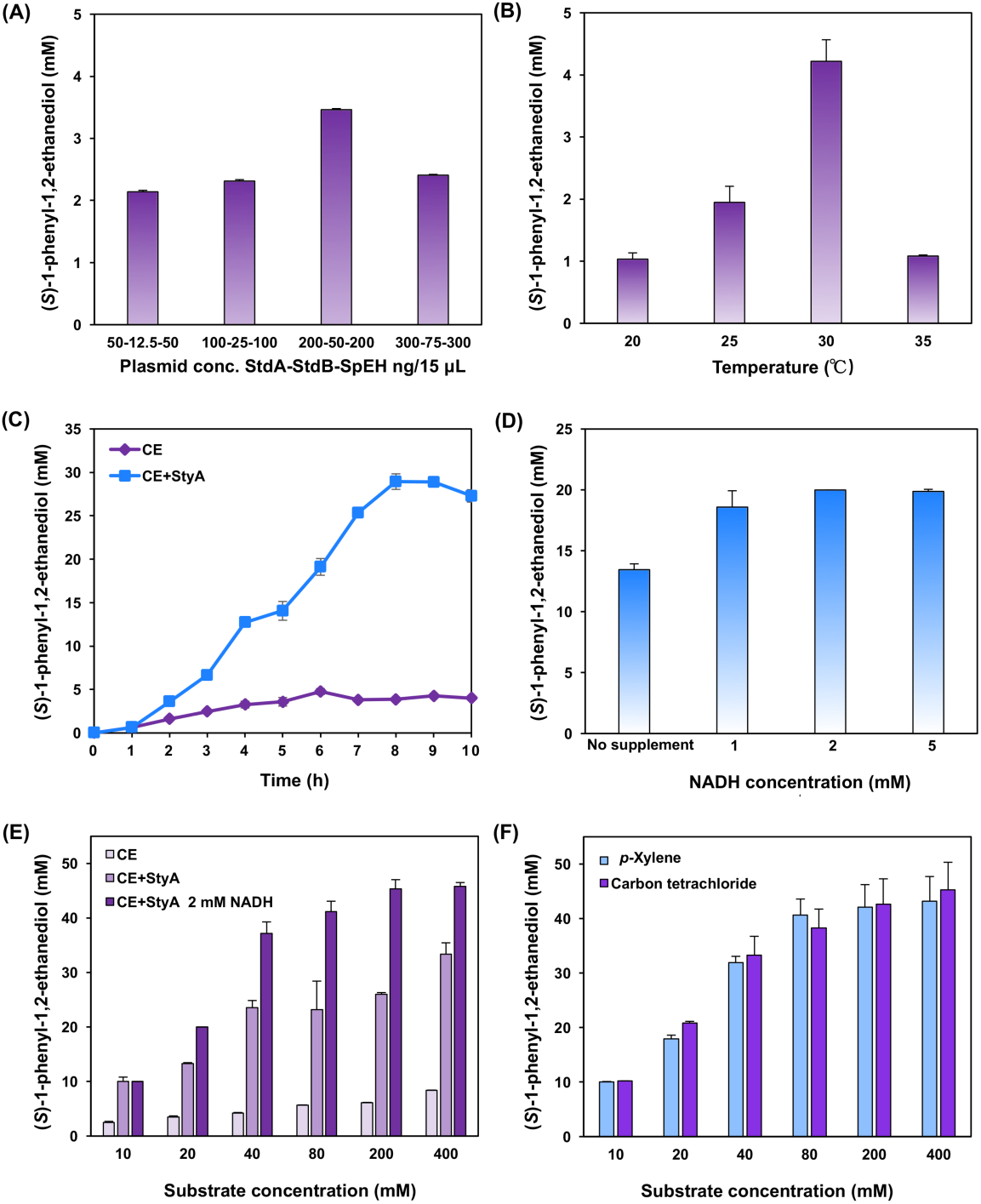
Optimization of (*S*)-1-phenyl-1,2-ethanediol production in the CFPS/emulsion system. **(A)** Optimization of plasmid concentration. Other conditions: substrate = 10 mM, reaction time = 10 h, reaction temperature = 30°C. **(B)** Optimization of reaction temperature. Other conditions: substrate = 10 mM, reaction time = 10 h, plasmid (StyA-StyB-SpEH) concentration = 200-50-200 ng/15 μL. **(C)** Time courses of (*S*)-1-phenyl-1,2-ethanediol production with 40 mM styrene. CE: cell extract without overexpressed StyA, CE+StyA: StyA-enriched cell extract. **(D)** Effect of additional supplement of NADH on (*S*)-1-phenyl-1,2-ethanediol formation with 20 mM styrene. **(E)** Production of (*S*)-1-phenyl-1,2-ethanediol with different styrene concentrations in three reaction systems. CE: cell extract without overexpressed StyA, CE+StyA: StyA-enriched cell extract, CE+StyA 2 mM NADH: StyA-enriched cell extract plus 2 mM NADH in the reaction. **(F)** Production of (*S*)-1-phenyl-1,2-ethanediol with different styrene concentrations in two CFPS/emulsion systems (*p*-xylene and carbon tetrachloride). Three independent CFPS reactions were performed for each condition, and one standard deviation is shown.

To test this hypothesis, we increased StyA in our biocatalytic system by overexpressing StyA in *E. coli* (see protein gel in **Figure S2**), followed by preparing StyA-enriched cell extracts to coexpress StyB and SpEH in CFPS reactions. By doing this, the conversion rate was notably increased up to 72% after an 8 h reaction with the (*S*)-1-phenyl-1,2-ethanediol concentration of 28.9 mM (**Figure 4C**). By contrast, without StyA-enriched cell extract in CFPS, the conversion rate was only about 10% in the presence of 40 mM styrene. Therefore, StyA-enriched cell extract was applied to our following CFPS reactions. Previous studies have shown that the activity of StyB depends on NADH consumption, which acts as an electron donor (Otto et al., 2004). We therefore next sought to supply additional NADH to the CFPS system and investigate the effect of NADH concentration on the product synthesis. As shown in **Figure 4D**, with the addition of NADH, the product yields obviously increased and 2 mM NADH gave rise to the highest yield of 20 mM (100% conversion based on 20 mM styrene). We next attempted to test a broad range of the substrate concentrations from 10 to 400 mM to see if our cell-free system was robust for the cascade biotransformation. Notably, 100% conversion was achieved with low substrate concentrations (<20 mM). However, further increase of styrene concentrations from 40 to 400 mM reduced conversion rates from 93% to 11%, respectively (**Figure 4E**, see the group of “CE+StyA 2 mM NADH”), perhaps as a result of toxicity and inhibition of styrene on the enzymes. Overall, in our current optimized cell-free system, we obtained the highest yield of approximately 45 mM, which is 113-fold higher than the initial yield (0.4 mM) of a control using only the aqueous phase (see **Figure S3** for initial data). Importantly, our cell-free system enables complete conversion (100%) of styrene at low concentrations (10 and 20 mM), which is higher than previous reported whole-cell biotransformation in either two-liquid-phase (92% conversion, 20 mM styrene) (Wu et al., 2014) or membrane-based biphasic (30% conversion, 20 mM styrene) systems (Gao et al., 2017). The higher conversion rate in our CFPS system is likely in part due to the faster mass transfer without cell walls or other membrane boundaries.

To demonstrate our system for potential applications, we were curious to know if CFPS might work with other organic solvents, since there might be some benefits being able to use different solvents to achieve other cascade biotransformations. We, therefore, investigated five other solvents (*i*.*e*., ethyl acetate, tetrahydrofuran, carbon tetrachloride, methylene chloride, and *p*-xylene) that can dissolve the copolymer PCL_2_1-PEG_45_-PCL_21_. We initially expressed sfGFP in different solvent systems. Our data indicate that sfGFP can be synthesized in high-yield (>1 mg/mL) from two out of the total five solvents, which are carbon tetrachloride and *p*-xylene (**Figure S4**). We next used these two organic solvents to prepare aqueous-organic emulsions for cascade reactions. As shown in **Figure 4F**, both systems worked like the toluene system that 100% conversion was observed with a low styrene concentration (10 mM), but it also gradually reduced to around 10% at 400 mM substrate. This success suggests that cell-free expression of many other enzymes, being able to catalyze broad organic soluble substrates, might be also possible in our emulsions. On the other hand, more choice of organic solvents demonstrates the robustness, flexibility, and applicability of our CFPS system in different aqueous-organic emulsions.

Finally, we scaled up our reaction system to 3 mL (1.5 mL CFPS + 1.5 mL emulsion) performed in a 50 mL tube. When using a moderate concentration of styrene (80 mM) as a starting substrate, we observed a yield of 21 mM (26% conversion). This is comparable to the small scale reaction (180 μL), which had a yield of 23 mM (29% conversion). Our consistent results between an order of magnitude increase in reaction scale suggests the possibility for using cell-free systems to meet the demand for future synthesis of chemicals. In addition, we investigated the reusability of our system and found that the relative activity of the third cycle still retained 62% of the initial activity (**Figure S5**). While our system can be reused, more work needs to be done in the future to keep the system’s high activity in terms of reusability.

In summary, we demonstrated the ability to use CFPS in a biphasic system, which we anticipate will provide a new, potential platform approach for chemists and biologists to synthesize valuable chemicals when water-insoluble substrates, bioconversion rates, mass transfer, and cellular toxicity limit the feasibility of whole-cell cultivation/fermentation. This platform has several key features. First, the enzymes necessary for the biotransformation were expressed by CFPS without purification and these enzymes performed biocatalytic reactions *in situ* at the aqueous/organic interface. Second, our optimized system was highly efficient, achieving 100% conversion of styrene to (*S*)-1-phenyl-1,2-ethanediol with substrate concentrations below 20 mM. Surprisingly, this system was only possible when StyA was enriched in the extract source strain by overexpression and StyB, which is insoluble when expressed in cells, was made by cell-free protein synthesis. Third, we discovered that the CFPS system was tolerant of a variety of organic solvents, including toluene, carbon tetrachloride, *p*-xylene, ethyl acetate, and methylene chloride. Fourth, we showed linear scalability of product titers and conversion yields over an order of magnitude range in volume. Fifth, we demonstrated the reusability of our system with several operational cycles. Taken together, these features show the robustness and scalability of the CFPS/emulsion system for biotransformation efforts. Looking forward, we envision that our cell-free system will enable new directions in developing biocatalytic cascades for defined biotransformations, providing a new and feasible avenue for efficiently synthesizing and manufacturing chemicals of pharmaceutical and industrial importance.

## Acknowledgments

The authors would like to acknowledge Ashty Karim for helpful discussions. This work was supported by grants from the National Natural Science Foundation of China (31971348 and 31800720), the Natural Science Foundation of Shanghai (19ZR1477200), and the Shanghai Pujiang Program (18PJ1408000). J.L. also thanks the starting grant of ShanghaiTech University. M.C.J. acknowledges support from the Department of Energy Grant DE-SC0018249, the David and Lucile Packard Foundation, and the Camille Dreyfus Teacher-Scholar Program.

## Supporting Information

### Materials and methods

#### Chemicals

Styrene, styrene oxide, (*S*)-1-phenyl-1,2-ethanediol, and toluene were purchased from Adamas (Shanghai, China). The triblock copolymers PCL_21_-PEG_45_-PCL_21_ of poly(ethylene glycol)-block-poly(*ε*-caprolactone) was prepared as described previously.^1^ The plasmid miniprep kit was purchased from Sangon biotech (Shanghai, China). All other chemical reagents were of highest purity available.

#### Strains, media, and plasmids

*E. coli* DH5α and BL21 Star (DE3) were used for plasmids propagation and cell extract preparation, respectively. The LB medium (10 g/L sodium chloride, 5 g/L yeast extract, and 10 g/L tryptone) was used for *E. coli* cultivation. The 2xYTPG medium (10 g/L yeast extract, 16 g/L tryptone, 5 g/L sodium chloride, 7 g/L potassium hydrogen phosphate, 3 g/L potassium dihydrogen phosphate, and 18 g/L glucose, pH 7.2) was used to grow cells for cell extract preparation. All genes used in this work were synthesized and cloned into plasmid pET28a by GenScript (Nanjing, China), including styrene monooxygenase (SMO, GenBank accession no.: AF031161) from *Pseudomonas* sp. VLB120 and epoxide hydrolase (SpEH, GenBank accession no.: KX146840) from *Sphingomonas* sp. HXN-200.

#### Preparation of cell extracts

Cell growth, collection, and extracts were prepared as described previously.^2^ Briefly, 1 L of 2xYTPG medium was inoculated with overnight preculture at an initial OD_600_ of 0.05. When the OD_600_ reached 0.6-0.8, cells were induced with 1 mM IPTG, followed by harvest at an OD_600_ of 3.0. Then, cells were washed three times by cold S30 Buffer (10 mM Tris-acetate, 14 mM magnesium acetate, and 60 mM potassium acetate). The pelleted cells were resuspended in S30 Buffer (1 mL/g of wet cell mass) and lysed by sonication (10 s on/off, 50% of amplitude, input energy ∼600 Joules). The lysate was then centrifuged twice at 12,000 g and 4°C for 10 min. The resulting supernatant was flash frozen in liquid nitrogen and stored at -80°C until use.

#### Cell-free protein synthesis (CFPS)

Standard CFPS reactions were carried out in 1.5 mL microcentrifuge tubes at 30°C for 10 h. Each reaction (15 μL) contains the following components: 12 mM magnesium glutamate, 10 mM ammonium glutamate, 130 mM potassium glutamate, 1.2 mM ATP, 0.85 mM each of GTP, UTP, and CTP, 34 μg/mL folinic acid, 170 μg/mL of *E. coli* tRNA mixture, 2 mM each of 20 standard amino acids, 0.33 mM nicotinamide adenine dinucleotide (NAD), 0.27 mM coenzyme A (CoA), 1.5 mM spermidine, 1 mM putrescine, 4 mM sodium oxalate, 33 mM phosphoenolpyruvate (PEP), 13.3 μg/mL plasmid, and 27% (v/v) of cell extract. The synthesized proteins were analyzed by SDS-PAGE and Western Blot. The concentration of sfGFP was calculated based on the fluorescence intensity detected by using the microplate reader (SYNETGY H1). To do this, after CFPS reactions, the emulsion was centrifuged at 12,000 g for 5 min to separate the aqueous and organic phases. The aqueous phase containing sfGFP was used for fluorescence measurement. Two microliters of the aqueous phase were mixed with 48 μL nuclease-free water and placed in a 96-well plate with flat bottom. Then, measurements of the sfGFP fluorescence were performed with excitation and emission wavelength at 485 and 528 nm, respectively. The fluorescence of sfGFP was converted to concentration (mg/mL) according to a linear standard curve made in house.

#### Cell-free reactions for cascade biotransformation

Cell-free reactions for (*S*)-1-phenyl-1,2-ethanediol production were performed in 2 mL microcentrifuge tubes at 30°C and 150 rpm for 10 h unless otherwise noted. The reaction emulsion (180 μL) contained CFPS mixture (aqueous phase) and PCL_21_-PEG_45_-PCL_21_ (organic phase, 20 mg/mL in toluene) at a volume ratio of 1:1. The substrates styrene and styrene oxide were dissolved in toluene and used at appropriate concentrations. For scale-up reactions with 3 mL emulsion, the reactions were carried out in 50 mL centrifuge tubes.

#### Analytical methods

CFPS mixtures were extracted by ethyl acetate for 4-6 times (as thoroughly as possible) and were redissolved in methanol. The gradient diluted (*S*)-1-phenyl-1,2-ethanediol were prepared in the same way as standards. The concentrations of styrene, styrene oxide, and (*S*)-1-phenyl-1,2-ethanediol were analyzed by using GC-MS. Helium was used as the carrier gas at a flow rate of 1.0 mL/min, and the detector temperature was 250°C. The column temperature was programmed from 60°C to 300°C at 30°C/min and hold at 300°C for 2 min. The retention times of styrene, styrene oxide, and (*S*)-1-phenyl-1,2-ethanediol were 3.21 min, 4.28 min, and min, respectively. All measurements were performed in triplicate.

## Supplementary figures

**Figure S1.**
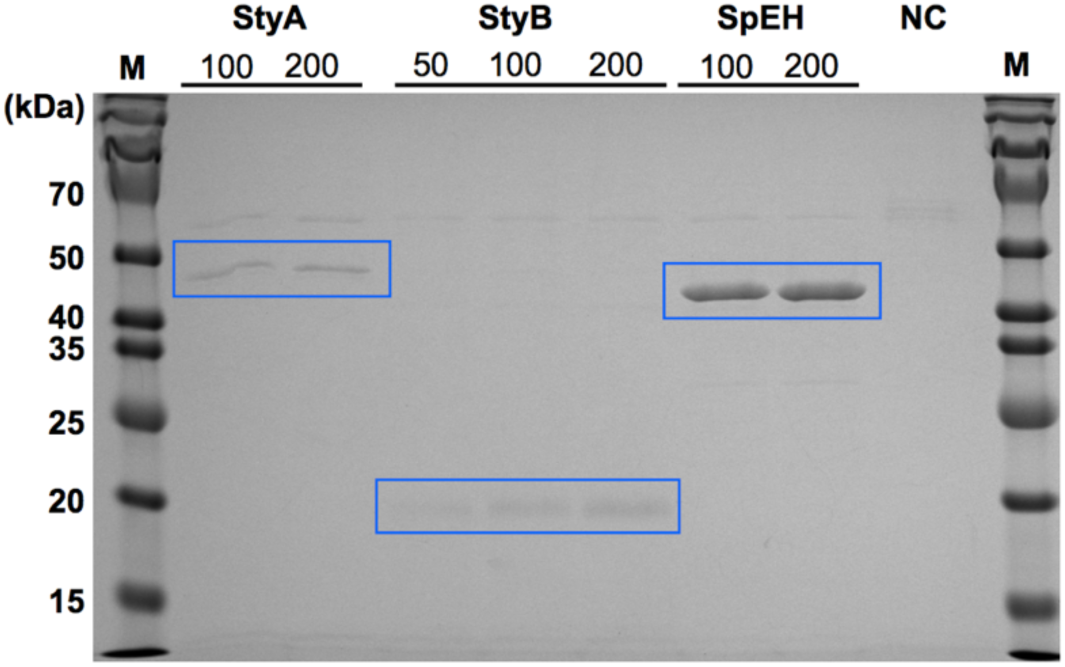
SDS-PAGE analysis of purified StyA (46.3 kDa), StyB (18.4 kDa), and SpEH (42.9 kDa) from CFPS reactions (each protein was purified from 270 μL (3 × 90 μL) reaction mixture). 50, 100, and 200 indicate plasmid concentrations (ng/15 μL) per reaction. NC, purification from CFPS reactions without plasmids.

**Figure S2.**
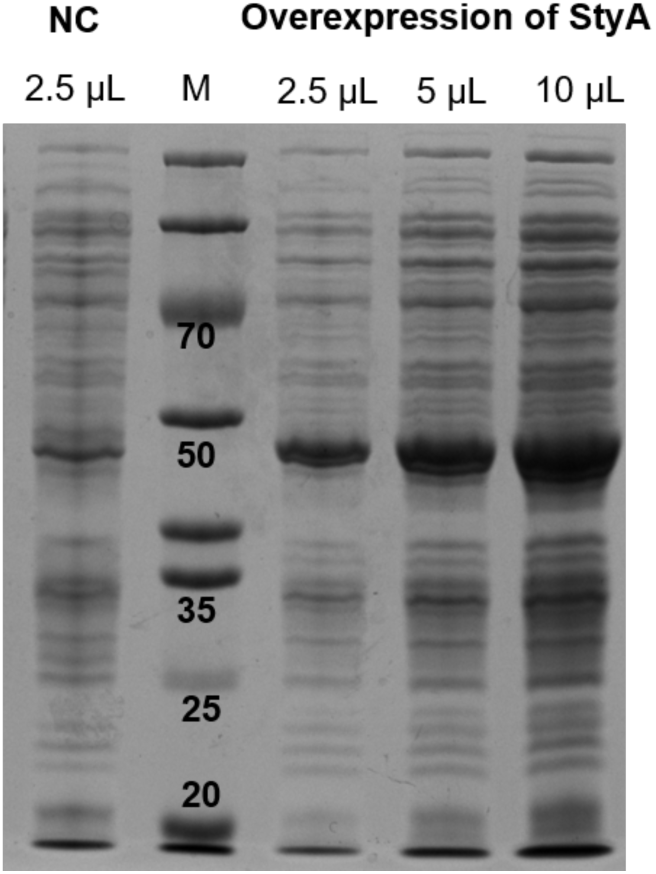
SDS-PAGE analysis of StyA (46.3 kDa) overexpressed in *E. coli* BL21 Star (DE3). 2.5, 5, and 10 μL indicate sample volume loaded to the gel. NC: negative control without overexpression of StyA. M: protein marker (kDa).

**Figure S3.**
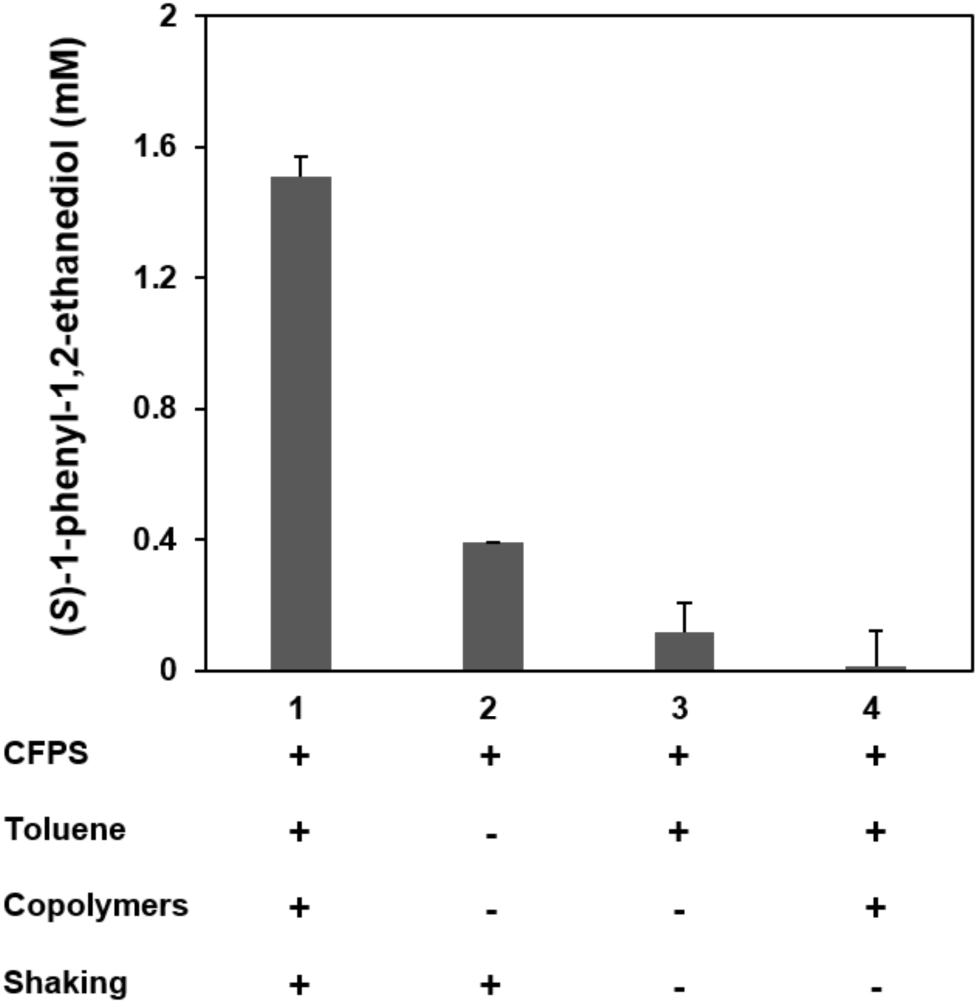
Yields of (*S*)-1-phenyl-1,2-ethanediol in different reaction systems. +: addition to the reaction. -: no addition to the reaction. Cell extract: *E. coli* BL21 Star (DE3). Three independent CFPS reactions were performed for each condition, and one standard deviation is shown.

**Figure S4.**
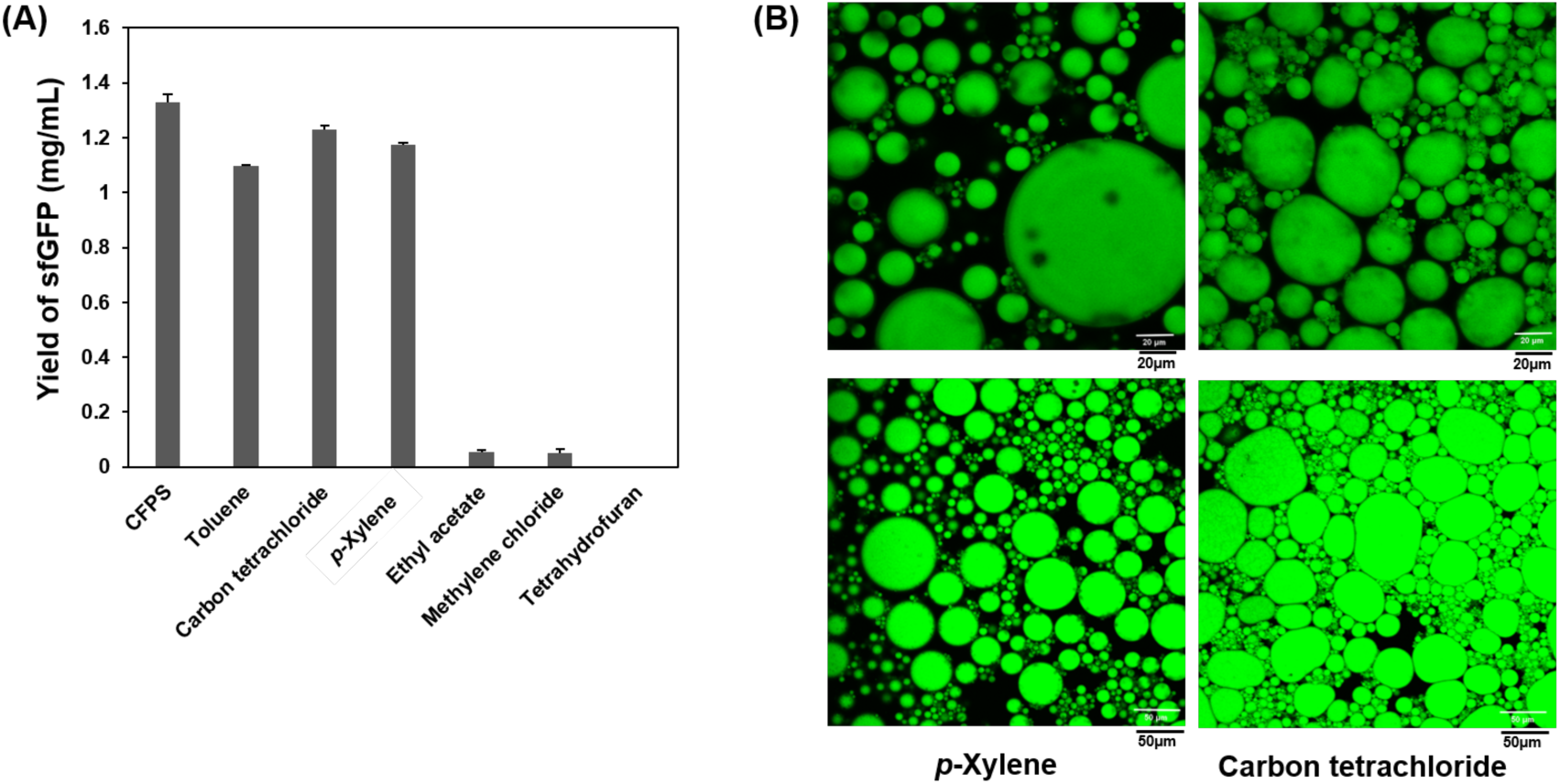
(A) Yields of sfGFP in different CFPS/emulsion systems. Three independent CFPS reactions were performed for each condition, and one standard deviation is shown. (B) Structures of PCL_21_-PEG_45_-PCL_21_ emulsion in *p*-xylene and carbon tetrachloride organic solvents detected by confocal laser scanning microscopy (CLSM) (top: 40x and bottom: 20x).

**Figure S5.**
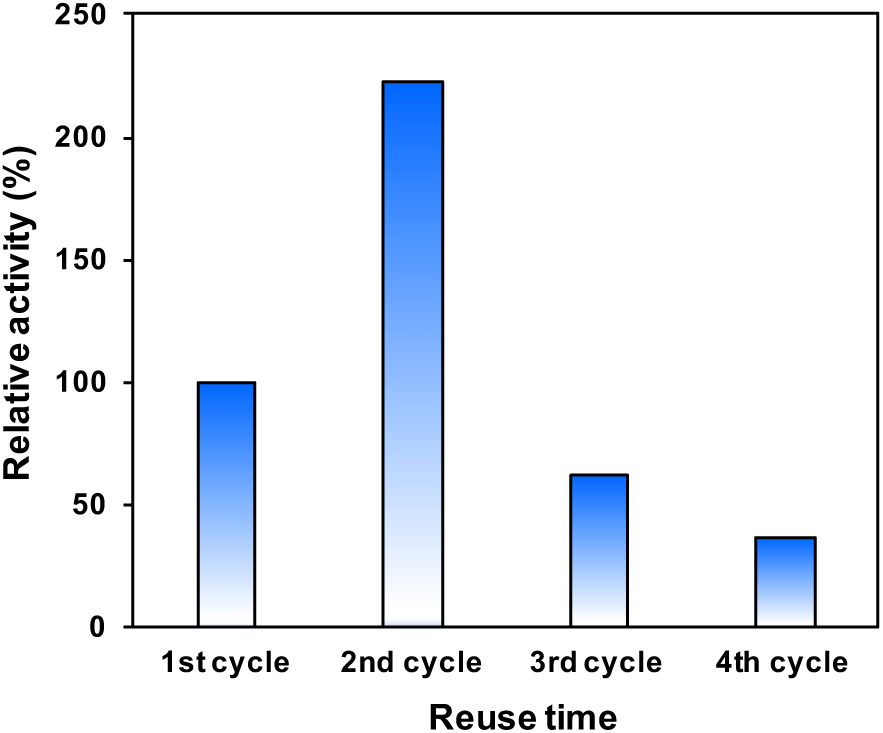
Reusability of the cascade reaction system. The system was reused four times and each cycle was performed for 3 h. At the end of each cycle, the organic phase was removed as more as possible and the fresh organic phase with substrate was added. The relative activity of the second cycle was more than 200% as compared to the first cycle. This is because enzymes need to be expressed during the initial hours and, therefore, in the second cycle more enzymes were expressed and showed a higher activity. The third and fourth cycles retained 62% and 37% of the initial activity, respectively.

